# A study’s got to know its limitations

**DOI:** 10.1101/2020.04.29.067843

**Authors:** Phil Gooch, Emma Warren-Jones

**Affiliations:** Scholarcy Limited

**Keywords:** meta-research, research rigour, study limitations, scholarly communication

## Abstract

**Background:** All research has room for improvement, but authors do not always clearly acknowledge the limitations of their work. In this brief report, we sought to identify the prevalence of limitations statements in the medRxiv COVID-19 SARS-CoV-2 dataset.

**Methods:** We combined automated methods with manual review to analyse manuscripts for the presence, or absence, either of a defined limitations section in the text, or as part of the general discussion.

**Results:** We identified a structured limitations statement in 28% of the manuscripts, and overall 52% contained at least one mention of a study limitation. Over one-third of manuscripts contained none of the terms that might typically be associated with reporting of limitations. Overall our method performed with precision of 0.97 and recall of 0.91.

**Conclusion:** The presence or absence of limitations statements can be identified with reasonable confidence using automated tools. We suggest that it might be beneficial to require a defined, structured statement about study limitations, either as part of the submission process, or clearly delineated within the manuscript.

## Introduction

All research has room for improvement, but authors do not always clearly and specifically acknowledge the limitations of their work [1][2]. Previous studies on the reporting of limitations in biomedical literature found that over 25% of 300 sampled papers contained no limitations [3]; while an earlier study, using automated methods to screen 400 articles for the presence of cue words, suggested that 83% did not mention any identifiable limitations statements [4].

To improve dissemination and reporting of the scholarly record, some have argued that journals should make authors’ study limitations public and freely available [5]. Typically, limitations are detailed in the Discussion section [6], although occasionally they may also appear in the Abstract [3]. To standardise recording of limitations, and to make a study’s shortcomings more transparent, some have argued that research papers should always have a separate, structured section dedicated to this [5].

Inspired by this previous work, and in particular by concerns raised about the rush to publish research on COVID-19/SARS-CoV-2 [2], we sought to answer the following questions about medical preprints on the novel coronavirus:

- What proportion contain a clear discussion of their limitations?
- How easy is it to find the discussion of the limitations?
- Where in the paper are the limitations mentioned?

Since 2019, medRxiv [7] has become the major repository for distribution of unpublished, non-peer reviewed literature on medical and health research. We decided to use manuscripts posted on medRxiv as the dataset for this brief study.

## Methods

We took the medRxiv feed of COVID-19 SARS-CoV-2 preprints from https://connect.medrxiv.org/relate/content/181 on 24 April 2020, which at the time listed 1747 manuscripts on the medRxiv server, and around 490 on the related bioRxiv server. We selected just the 1747 medRxiv manuscripts.

These manuscripts had been made available on the medRxiv server between 10 January 2020 and 22 April 2020. All manuscripts were PDF files, ranging in size from < 1MB to over 35MB. Following [4], our aim was to use automated methods to analyse the manuscripts as far as possible, but also follow [3] by cross-checking the automated results with manual inspection and verification.

While these medRxiv manuscripts are also available in machine-readable format as part of the AllenAI CORD-19 dataset [8], the full-text in that dataset is in the form of body text chunks delimited by citation spans, and would have required additional pre-processing to add the structure we required (see Discussion). Instead, the manuscripts were processed through the Scholarcy API^1^ to convert the PDFs to structured data in JSON format, which includes the full text of the manuscript, structured into the main section headings and their content. The API makes use of regular-expression style pattern matchers over spacy.io^2^ linguistic features to identify section headings. It also uses a sentence-level statistical classifier, trained on character n-gram features from the PubMed OpenAccess Subset^3^ to identify funding statements, ethical compliance statements, data availability statements, and limitations statements.

From the API output, we identified how many manuscripts recorded limitations in a structured way in a specific section/subsection by counting the number of section headings across the corpus containing the word ‘Limitations’. We also used the output of the API’s limitations statement classifier to identify how many manuscripts contained limitations statements generally in the text, and in which section these appeared. We then checked the results by hand to calculate the precision and recall of this automated process.

## Results

The section-heading extraction identified 516 ‘limitations’-type headings, subheadings, or run-in headings/paragraph starts in 494 of the 1747 manuscripts (28%). Manual inspection revealed 22 false positives, although all 494 manuscripts contained at least 1 correct limitations statement. Removal of the 22 false positives give a precision of 1-22/516 = 0.96.

The limitations classifier identified 930 limitations statements in 834 of the 1747 manuscripts (48%). Manual inspection revealed that 28 of the statements automatically classified as a limitation were false positives, which gives us a precision score of 1-28/930 = 0.97.

We reviewed the remaining 913 manuscripts to see how many of these did in fact contain limitations mentions that were missed by the automated extraction process. Of these 913 manuscripts, we found, by manual inspection, a discussion of limitations in only 83, which gives us a combined limitations headings extraction + limitations classifier recall of 1 - 83/913 = 0.91.

Adding these 83 manuscripts missed by the classifier to the 834 correctly flagged by the classifier gives us a total of 917/1747 = 52% of the dataset contained a verified limitations statement.

Table 1 provides a breakdown of the location of limitations statements in these 917 manuscripts. In 11 manuscripts, limitations were also restated in the abstract.

**Table 1:**
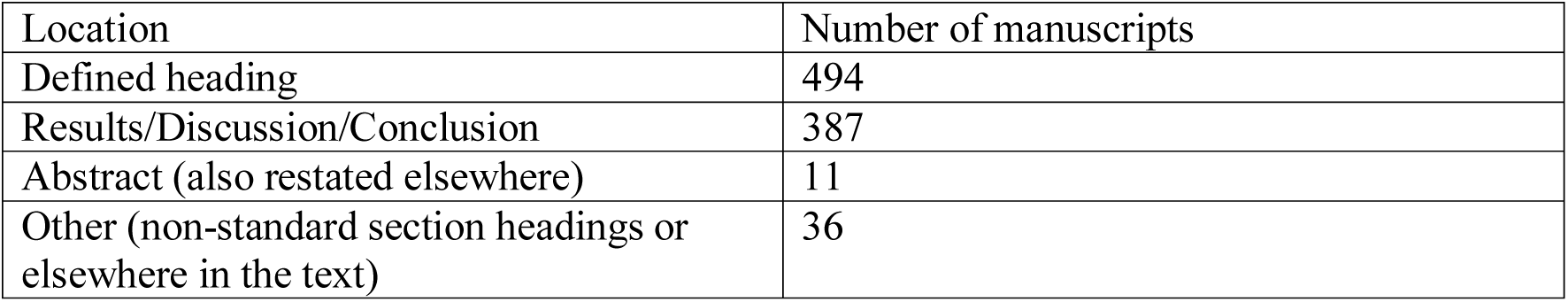
Where are limitations statements located?

Table 2 shows a representative sample of limitations statements missed by the classifier.

**Table 2:** Examples of limitations not identified by automated processing.

|  |
| --- |
| We conclude with a word of caution |
| We are certainly no experts in this field and a GM is simply a description of a smooth time evolution of infections |
| We leave it to the reader to treat the here presented observations and claims with enough care |
| We recognize that our data source is useful and timely but should not replace official statistics |
| One significant factor that could invalidate the predictions |
| Our results should be interpreted in relative terms |
| Our findings are constrained to a limited time frame and might have missed time-changing disease accounting for the limited follow-up duration |
| This study is a cross-sectional study, which only described and analyzed COVID-19 patients with positive real-time RT-PCR results |
| Take the exact figures obtained with our simulations with a grain of salt |
| Our results are based on data for a single, large country |
| Caveats: these are preliminary results |
| We caution that the data are sparse |
| Reported data may not be accurate |
| Our model should be used with extreme caution |
| The results do strongly depend on limitations on data collection |
| Due to shortcomings in the reported data |
| A word of caution concerning the present view: any model is but an imperfect view of reality |

It seemed surprising to us that 1747 - 913 = 830 manuscripts (48%) appeared to contain no clear mention of limitations, so as a sanity check, we ran a search across these to see if we had missed anything obvious. We searched for any combination of one or more of the words:

> *limitations, limitation, shortcomings, shortcoming, caution, defects, defect, weaknesses, weakness, deficiencies, deficiency, future research, future work, future studies, future study, further research, further work, further studies, further study*

across the full text within the JSON data. Yet 616 manuscripts (35%) contained none of these words.

## Discussion

In the medRxiv COVID-19 SARS-CoV-2 snapshot, we found limitations statements in 917 of the 1747 preprints - just over 52%. While undoubtedly some of the remainder probably did contain limitations statements, but we failed to find them, the fact that 35% of the total contained none of the words you might expect to appear at least once, somewhere in the paper, suggests that our results align with the 2013 study by Ter Riet et al [3]. In contrast to an earlier study [4], we manually checked the output for each manuscript, so our results are perhaps more realistic than the somewhat alarming 83% figure provided in [4].

It would be interesting to repeat this analysis over the larger CORD-19 dataset [8]. This dataset contains a combined bioRxiv/medRxiv subset which, at the time of writing, contained 2,278 JSON files^4^. In fact, a quick scan of this dataset reveals a similar paucity of defined limitations sections as found in the analysis of our own automated processing: searching the CORD-19 subset using the regular expression (case-insensitive):

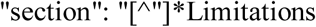

reveals matches in only 106 of the 2278 manuscripts (~5%), and 950 (42%) contain none of the sanity-check word-search terms.

The precision and recall of our automated limitations-statement classifier (0.97, 0.91, respectively; F1=0.94), which combines a pattern-based approach with machine learning, is similar to that of the tool developed and reported by [9]. It appears that, if a limitations statement is present, it should be reasonably straightforward to identify it in the text using an automated approach. Future work should verify whether these results hold for a much larger dataset of preprints (or published papers), more representative of a wider field of study, than the COVID-19 dataset considered here.

The dataset comprised unreviewed preprints that represent preliminary, breaking research. Those preprints that go on to journal submission and peer review are likely to go through a number of revisions that may make the reporting of limitations more explicit and clearly defined. But if preliminary research is to be put in the public domain, is it enough to add a ‘treat with caution’ disclaimer? Shouldn’t it be good practice, even with the need to accelerate publication during a crisis, for authors to lay out study limitations clearly and specifically, particularly as preprint findings start to be reported in the mainstream media?

Whether limitations belong in their own, labelled (sub)section is a decision for reviewers, journal editors and the wider research community. However, for someone screening preprints, it might make the job easier if these are clearly identifiable - almost as metadata - in the same way that other information such as funding, acknowledgements, data availability and conflicts of interests tend to be declared in a manuscript.

## Limitations

Only one author checked the automated identification of limitations statements extraction. The author is not a domain expert and so may not have picked up all limitations where they were not clearly expressed. Also, presence or absence of a limitations statement was checked against document text contained within the machine-extracted JSON data, but not against original manuscript PDF. It is possible that in some cases, the text extraction may not have extracted the complete article content. However, this risk was mitigated by performing post-hoc checks against the CORD-19 dataset [8]. Finally, the sanity-check search may not have been comprehensive, and limitations may have been stated in a more implicit way that required domain knowledge to interpret. As a result, the recall figure of the automated process may be lower, and the proportion of manuscripts in this collection that do in fact state their limitations may be higher than that reported here. Close reading of the original manuscripts by multiple domain experts would reveal a more accurate estimate of the prevalence of limitations statements, similar to [9].

## Conclusion

We used automated processing combined with manual review to estimate the prevalence of clearly identifiable limitations statements in the medRxiv corpus of COVID-19 SARS-CoV-2 preprints. We found that around half of them did not appear to contain such a statement, and just over a third of them contained none of the words that might be associated with reporting of limitations. We suggest that it might be beneficial to require a defined, structured statement about study limitations, either as part of the submission process, or clearly delineated within the manuscript.

## Data availability

The 24 April 2020 medRxiv COVID-19 dataset of 1747 PDF URLs, and the corresponding API output for each PDF, is available from the authors. As the copyright of the full text of each manuscript is with each author, and our analysis required full-text PDF extraction, we are unable to make the extracted full-text data publicly available. However, the Scholarcy API is available for those who wish to re-run the extraction from the list of URLs^5^, and machine-readable versions of these papers, albeit in a different format, are available as part of the CORD-19 Dataset [8].

## Acknowledgement

We thank J. Brian Byrd, MD, MS, for feedback on an earlier version of this report.

## Footnotes

1 https://api.scholarcy.com

2 https://spacy.io/usage/linguistic-features

3 https://www.ncbi.nlm.nih.gov/pmc/tools/openftlist/

4 https://ai2-semanticscholar-cord-19.s3-us-west-2.amazonaws.com/latest/biorxiv_medrxiv.tar.gz

5 https://api.scholarcy.com.

